# Interspecies bacterial competition determines community assembly in the *C. elegans* intestine

**DOI:** 10.1101/535633

**Authors:** Anthony Ortiz Lopez, Nicole M. Vega, Jeff Gore

## Abstract

From insects to mammals, a large variety of animals hold in their intestines complex bacterial communities that play an important role in health and disease. However, the complexity of these gut microbiomes and their hosts often constrains our ability to understand how these bacterial communities assemble and function. In order to elucidate basic principles of community assembly in a host intestine, we study the assembly of the microbiome of *Caenorhabditis elegans* with a bottom-up approach. We colonize the gut of the worm *C. elegans* with 11 bacterial species individually, in all possible pairs, and in selected trios, and we find an organized mixture of coexistence and competitive exclusion that indicates a hierarchical structure in the bacterial interactions. The capacity of a bacterial species fed in monoculture to colonize the *C. elegans* intestine correlates with its average fraction in co-culture experiments, yet fails to predict its abundance in many two- and three-species microbiomes. Hence, the bacterial fractional abundances in co-culture experiments—pairwise outcomes—are influenced by interspecies interactions. These pairwise outcomes accurately predict the trio outcomes in the worm intestine, further highlighting the importance of pairwise interactions in determining community composition. We also find that the *C. elegans* gut environment influences the outcome of co-culture experiments, and demonstrate that the low intestinal pH is one of the causes. These results highlight that a bottom-up approach to microbiome community assembly may provide valuable insight into the structure and composition of complex microbial communities.

## Introduction

Bacterial communities are found almost everywhere in nature. Among the many ecosystems in which bacterial communities play a fundamental role, the animal digestive tract is one of remarkable importance. These large, complex, and highly organized consortia can degrade food and deliver nutrients to their host, protect against invading pathogens, and even produce neurotransmitters that affect host behavior (1–7).

Despite considerable recent efforts toward elucidating the contents of these bacterial communities (8–11), the rules that govern their assembly are still poorly understood. Complicating this problem is the fact that these microbial communities are exceedingly complex and difficult to manipulate experimentally. The human gut microbiome, for instance, consists of approximately 4×10^13^ cells distributed across hundreds of species interacting in myriad ways across different regions of the gut (12). Moreover, these complex microbial communities vary between individuals based on many factors, including environmental conditions, diet, and the host genotype and immune system (13,14). This complexity and variability has stymied efforts to uncover general rules for microbial community assembly in the context of the host environment.

Current efforts have taken advantage of model systems to experimentally address the composition and assembly of simpler gut microbiomes (15,16). Animals such as zebrafish (17–19), honey bees (20–23), leeches (24–27), flies (28–31), and worms (32–35) can be raised and manipulated in the laboratory in order to assess their microbiomes. The bacterial communities associated with these animals contain a small number of taxa relative to humans, allowing bottom-up assembly of comprehensive, synthetic microbial communities. These experimentally tractable models provide an opportunity to observe population dynamics in host-associated microbiomes, by exposing a selected population of hosts to colonization under controlled conditions and monitoring the ensuing interactions among microbes and between microbes and host.

Here we use the nematode *Caenorhabditis elegans* (36) as a simple gut model to address issues of experimental tractability and to gain intuition about community assembly in microbiomes. Despite comprising only 20 epithelial cells, the *C. elegans* intestinal tract shares many physiological features with the intestines of higher organisms, including microvilli, a mucin layer, epithelial junctions, cyclical movement via contraction of muscles, and extensive interaction with the host immune system (37–43). The gut of *C. elegans* is well known to host stable populations of bacteria, including important pathogens and probiotics of humans, at substantial population levels up to 10^5^ bacterial cells per worm (34,44–46). Furthermore, recent reports have suggested that the gut of this millimetric organism is capable of filtering its bacterial environment and selecting a core microbiota (35).

In this study, we colonized the intestine of *C. elegans* with simple microbiomes to determine patterns in the assembly of microbial communities in a host intestine. We found that the inherent ability of a bacterial species to colonize the worm gut was an acceptable predictor of that strain’s average performance in two-species microbiomes, but these single-species data became inadequate at predicting the outcomes of specific two- and three-species communities. This suggests that interactions between bacterial species determine the shape and form of the microbial communities in this host. Importantly, we find that the outcome of two-species feeding experiments can be used to predict the composition of the three-species feeding experiments, indicating that “assembly rules” may provide insight into the composition of microbiome communities. Further, we identified a conserved bacterial competitive hierarchy between *in vivo* gut and *in vitro* liquid medium that is disabled in specific cases by the acidity of the *C. elegans* intestine. With this, we advance our understanding of the environmental filtering imposed by *C. elegans* and provide insight into bacterial community assembly.

## Results

To investigate the colonization and growth dynamics of different bacterial species in the gut of *C. elegans*, we fed germ-free synchronized adult worms with eleven single bacterial species over four days in a well-mixed liquid medium (Fig 1A, Methods). This length of feeding was long enough to ensure reliable colonization of the worms, yet short enough to avoid problems associated with pathogen-based killing by our strains (47). After this feeding and colonization period, we allowed worms to feed briefly on heat-killed bacteria to remove transient colonizers, cleaned the surface of the worms, and measured the intestinal bacterial densities by grinding batches of worms and counting colony forming units (CFU, hereafter called *cells*).

**Figure 1.**
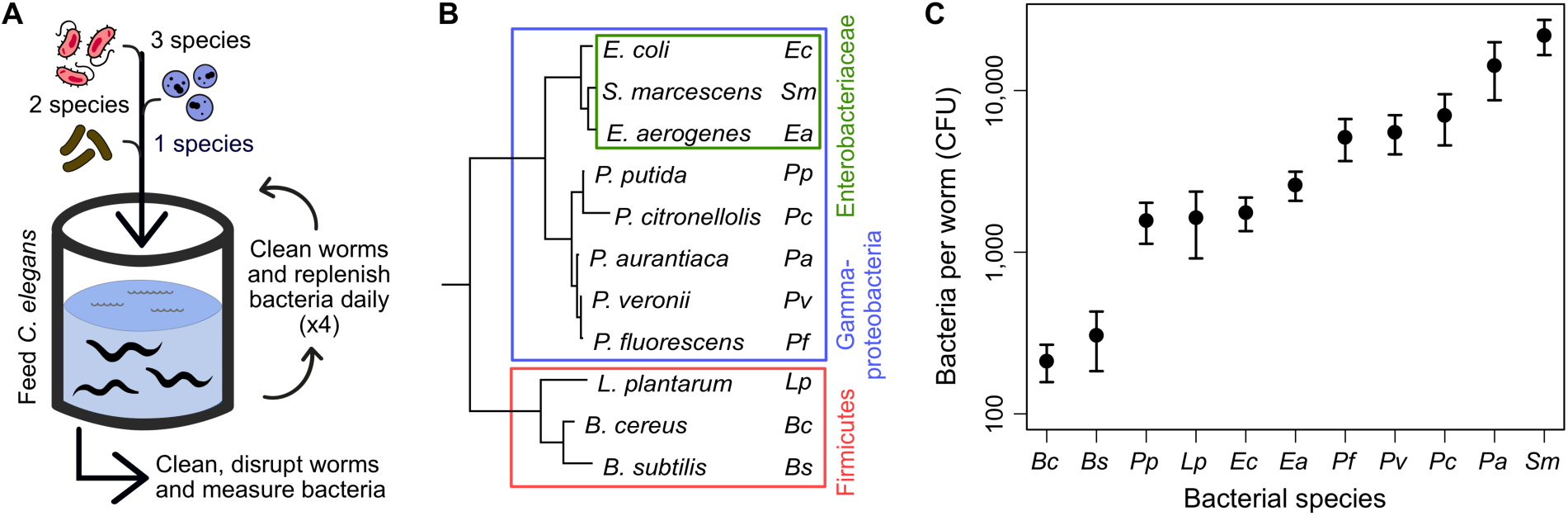
Eleven bacterial species colonize the gut of the worm in monoculture. **(A)** One, two or three bacterial species are fed in liquid culture to a sterile population of *C. elegans* AU37. Equal bacterial concentrations are maintained during 4 days of colonization. Afterwards, worms are mechanically disrupted in batches of ∼20 and CFU counts are used to determine bacterial population sizes. **(B)** Phylogeny of the 11 bacterial species used to colonize the gut of *C. elegans*. **(C)** Population sizes in monoculture colonization span two orders of magnitude and reflect the inherent capabilities of bacteria to colonize the worm intestine environment. Points represent the average of 8 or more biological replicates, and error bars as the standard error of the mean (SEM).

We utilized a set of eleven bacterial species representing the phyla Firmicutes (gram-positive) and Proteobacteria (gram-negative) (Fig 1B). From the latter phylum, we included soil isolates from the families *Enterobacteriaceae* and *Pseudomonadaceae*, abundant bacterial clades found in the core microbiota of *C. elegans* grown on its native environment of decaying organic matter (46,48,49), but none of our strains was directly isolated from nematodes. These experiments can therefore be viewed as probing community assembly of a gut microbiome with bacterial species encountered in the natural environment and without a period of evolutionary adaptation. We found that all bacterial species successfully colonized the *C. elegans* intestine, with population sizes ranging from two hundred cells per worm in the case of *Bacillus cereus* (*Bc*), up to tens of thousands of cells in the case of *Serratia marcescens* (*Sm*) (Fig 1C). These differences in bacterial population sizes in monoculture colonization reflect differences in the ability of each bacterial species to not trigger an aversive olfactory response in *C. elegans* (50–52), survive passage through the *C. elegans* grinder (53–56), adhere to the intestinal lumen (32,57–59), proliferate within the worm, or a combination of these features (60).

Despite the importance of revealing which of these characteristics allows each bacterial species to surpass the environmental filtering of the worm gut and reach a given population size, in this study we aimed to understand the influence of monoculture colonization in the assembly of the *C. elegans* microbiome. Thus we constructed the simplest intestinal communities by performing all possible co-culture experiments with these eleven bacterial strains (55 pairs in total). We fed worms in a well-mixed liquid medium with pairs of bacteria present at equal concentrations (*cells*/mL) to ensure that strains had the same initial probability to be ingested (Methods). We found that a majority (41/55∼75%) of pairs displayed coexistence of the two species (Fig 2A, left panels), whereas the remainder (14/55) led to competitive exclusion of a species (Fig 2B). Given that we are feeding the worms at equal concentrations, we cannot detect the possibility of bistability, in which the outcome of competition depends upon the starting ratio of the two species. Some bacterial species consistently exclude other species (*Sm*) or are excluded (*Bs*), but the large amount of coexistence (Fig S1) makes the simple winner/loser classification inappropriate for the observed bacterial competitive abilities.

**Figure 2.**
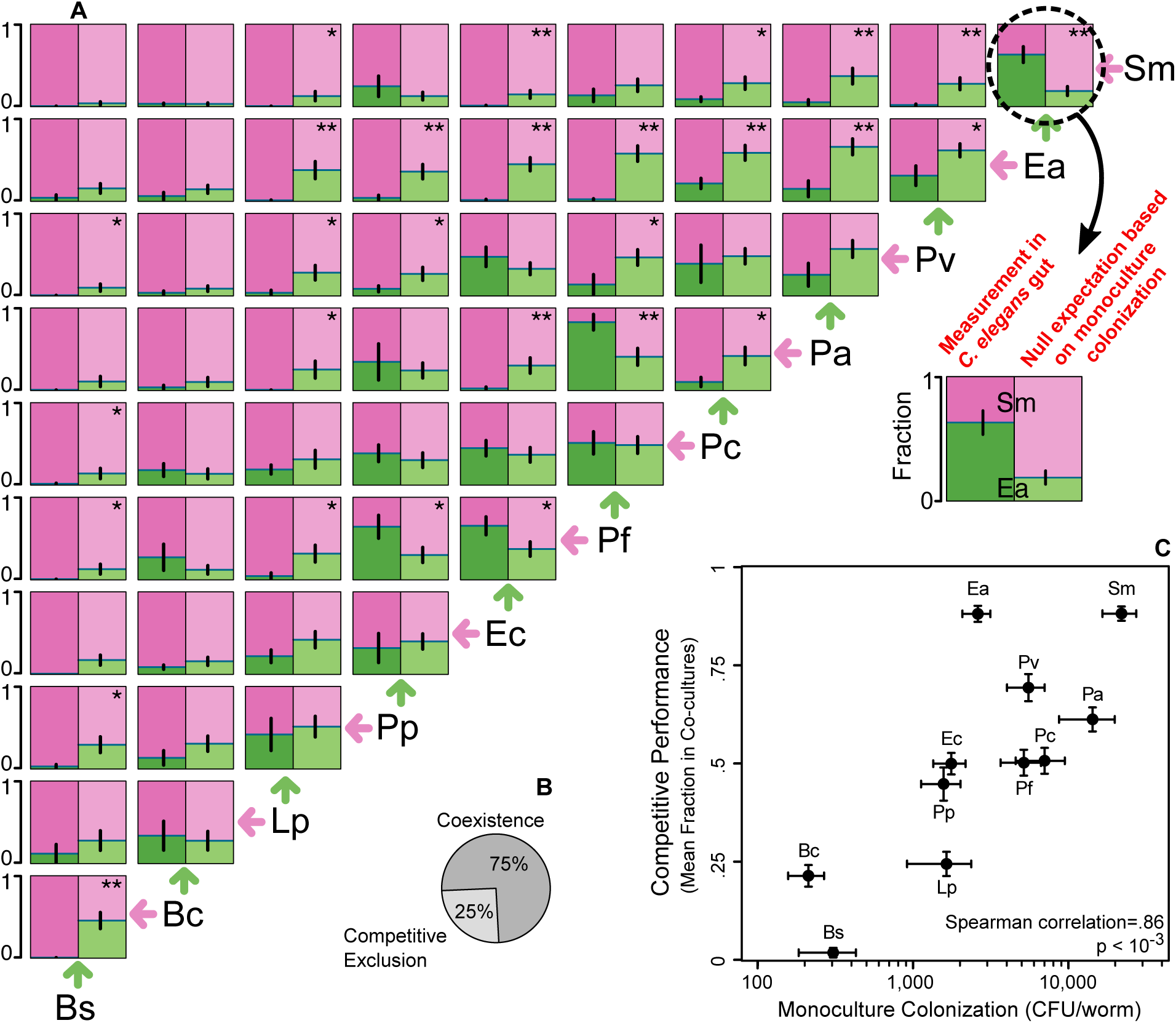
Monoculture colonization relates to, but does not predict pairwise outcomes in the worm intestine. **(A) Left:** Fractional abundances of 55 co-culture experiments inside the *C. elegans* intestine. Error bars as the SEM of 2 to 8 biological replicates (Fig S1). Bacterial strains are ordered by its mean fractional abundance. **Right:** Null expectation for the fractional abundances based on a non-interacting model where each bacterial species reaches its monoculture population size. Null expectation as the mean of all possible pairings of monoculture information, and error bars as the SEM (Methods). * and ** represent a statistically significant difference between measurement and null expectation at *p*-values of .05 and .01, respectively (Welch’s T-test). **(B)** Coexistence is common and competitive exclusion at a detection limit of 2% is rare. **(C)** Competitive performance, as the mean fractional abundance in co-culture experiments, correlates with monoculture population size. X-axis as in Fig 1B, and error bars on Y-axis as the propagated error from the SEM of each pairwise competition.

A better summary-metric of the pairwise outcomes is the mean fractional abundance of a species when competed against each of all the other species. Full dominance, extinction, and coexistence award 1 point, 0 points, and the fraction of point at which the species is present in each co-culture experiment, respectively, and the sum is then divided by the number of competitors. We found that this competitive performance score correlates to the population size reached in monoculture colonization (Fig 2C, Spearman correlation [r_s_]=.86, *p*<10^-3^). This positive relationship indicates that a bacterial species persists in two-species microbiomes due to similar properties to those favoring its monoculture colonization of the gut. Interestingly, the population sizes do not correlate to the mean relative yields (*cells*/worm of a species in competition divided by *cells*/worm of the same species in monoculture), indicating that a large population size does not protect a bacterial species from being harmed by competition in co-culture experiments (Fig S2).

The interactions we measured can be categorized as hierarchical, since a highly ranked competitor will frequently exclude or dominate a low-rank adversary (dominance of pink-color in left panels of Fig 2A). The hierarchy score of this network, 0.82, is significantly larger than the hierarchy score found in random matrices with the same distribution of pairwise outcomes (*p* <10^-5^, Methods), suggesting that the hierarchy seen in our experiments is caused by bacterial features acting in a transitive manner. Consistent with this, we do not observe any cases of intransitive competition, in which the pairwise interactions of three bacterial species would be analogous to the rock-paper-scissors game and no absolute winner would exist (61,62). This intransitivity has been proposed as a major mechanism inducing coexistence in natural populations (63–67), but we do not observe it in any of our 165 possible trios. With a more relaxed definition of non-transitivity, in which a species wins a competition by being more abundant than the competitor instead of needing to fully exclude it, we find two candidate trios with a rock-paper-scissors-like structure: *Ec-Pf-Pa* and *Pp-Pf-Pa* (although the dominance of some competitors is not statistically significant).

*Lp, Bs*, and *Bc*, the three species representing the Firmicutes phylum, were competitively excluded in most co-culture experiments, consistent with their poor colonization in monoculture. Likewise, the strongest single species colonizer, *Sm*, dominates the final population in most of its pairwise competitions. However, not all strains and interactions acted according to expectations from single species colonization (68–70). For example, even though its monoculture population size was just above average, *Enterobacter aerogenes* (*Ea*) was capable of outcompeting most of its adversaries, suggesting that direct interactions between microbial species can play a significant role in determining the outcome of pairwise colonization.

We therefore sought to determine whether interactions between microbes are an important determinant of community structure, or whether monoculture colonization ability is sufficient to predict the outcomes of pairwise competition in the host. We calculated a null expectation for the fractional abundances assuming that each species is able to reach the carrying capacity that was measured in monoculture colonization (Fig 2A, right panels). By comparing this null expectation with the experimentally measured fractions obtained from pairwise colonization (Fig 2A, left panels), we are able to identify the cases in which interspecies interactions play a dominant role in determining the composition of the gut microbiome. Across all pairs, the measured fractions deviate from the null expectations by a mean distance of .2, with a peak at low distance and a long tail corresponding to cases where the interspecies interactions are particularly important (Fig S3). In 28 cases this deviation is large enough to reject the null model (*p*<.05), many more than the 2.75 cases expected by chance at this significance level and with these 55 pairwise combinations. This result indicates that a null expectation, where each species’ abundance in pairwise colonization is largely determined by the initial filtering event rather than by interactions between bacteria, is valid for some but not all pairs of species in this set.

The importance of interspecies interactions is further supported by an analysis of relative yields (Fig S4). Most species cannot reach their monoculture population size in co-culture experiments (relative yield<1), suggesting that interactions between species are largely competitive. The 21 out of 110 cases (19%) where we measured a relative yield larger than one could be due to modest facilitation or due to variation in colonization and measurement between the monoculture and pairwise experiments. Collectively, these data suggest that the monoculture colonization ability of a bacteria relates to its abundance in pairwise experiments, but the mostly competitive interspecies interactions alter as many as half of the individual pairwise outcomes (Fig 2, S4).

### Environmental filtering imposed by C. elegans

Next, we sought to characterize the environmental filtering performed by *C. elegans* during microbial community assembly. Specifically, we sought to quantify the differences in bacterial relative abundances between the *in vivo* worm gut and the *in vitro* liquid medium used as feeding substrate (Fig 3). Thus, we performed all fifty five co-culture experiments in liquid medium and measured the equilibrated bacterial fractions after seven cycles of daily dilution (Methods, Fig S5). We found that competitive performance in the worm gut and liquid media are correlated (Fig 3A), which suggests that the environmental filtering that *C. elegans* provides is not strong enough to alter the hierarchical order of these eleven bacterial strains. This resemblance between the worm gut and liquid media with the coarse-grained mean fractional abundance is also observable when comparing the mean relative yields (Fig 3A, inset).

**Figure 3.**
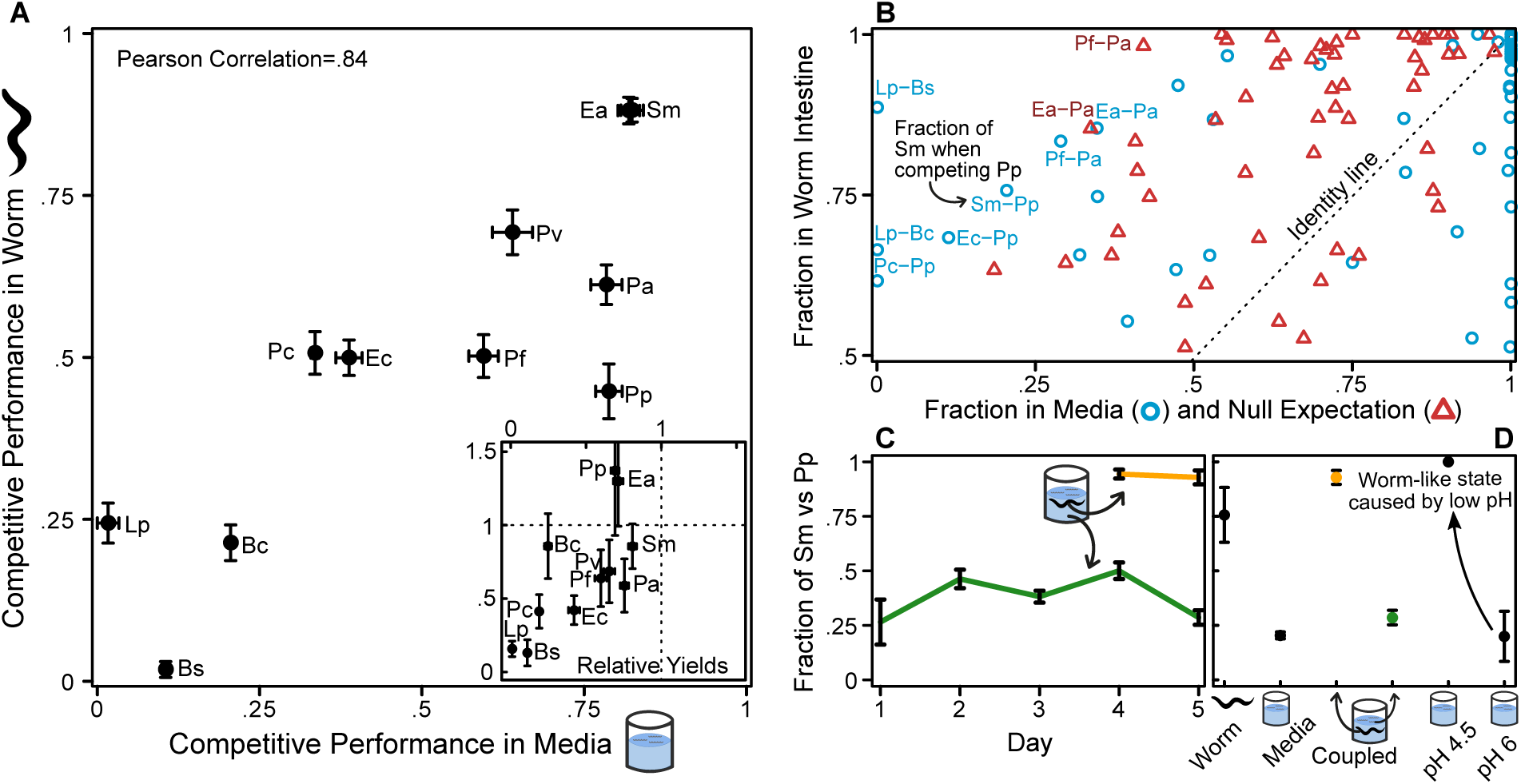
Bacterial competitive hierarchy is conserved between *in vivo* gut and *in vitro* liquid media environments. **(A)** Mean fractional abundance in co-culture experiments in the worm and in the background feeding media correlate with each other, indicating that strong competitors remain strong, and weak competitors remain weak in these two environments. **(Inset)** Mean relative yields (*cells*/worm of a species in competition divided by *cells*/worm in monoculture) in the worm and media also correlate. Error bars as the propagated error from the SEM of co-culture experiments. **(B)** Fractional abundances in co-culture experiments in worm intestine are as different from the background media as from the null expectation based on monoculture colonization. Points at distance >.5 from the identity line were labeled. **(C)** Fraction of *Sm* in worm (yellow) and media (green) when environments are coupled by migration. Both environments reach their different equilibrium points in the same test tube. **(D)** An acidic version of the media resembling the average pH of the worm intestine (4.5) shifts back the pairwise outcome of *Sm-Pp* to a worm-like state. Error bars in C and D as the SEM of at least 4 replicates.

Although competitive performance is similar between the worm gut and *in vitro* liquid media, we find that individual pairwise outcomes often display very different outcomes in these two environments (Fig 3B). We quantify the competitive performance of a bacterial species by averaging over all ten pairwise experiments; although these averages correlate between the worm intestine and the background media (Fig 3A), the underlying pairwise outcomes can be very different (Fig 3B, blue circles). The mean distance between the fractions in the media and the worm (0.22) is similar to the mean distance between the null expectation from monoculture colonization and the worm outcomes described earlier (Fig 3B, red triangles; Fig S3). If we wanted to predict the relative abundance of two-species communities in the worm intestine, both the monoculture colonization and the media pairwise outcomes would fail to a similar extent, but in dramatically different ways: while the null expectation overestimates the frequency of coexistence in the gut, the media competition displays less coexistence than observed in the worm gut. Our data therefore suggests that the *C. elegans* intestine doesn’t alter the competitive hierarchy of its bacterial colonizers, but is capable of shifting specific pairwise outcomes.

In fact, several pairs of species displayed remarkably different outcomes in competition inside versus outside of the worm intestine, such as *Sm-Pp* and *Lp-Bs*. Since the two environments were capable of selecting a different microbial community composition, we wondered if migration between them would homogenize their microbial compositions. To answer this question, we fed again a population of worms with a mixture of *Sm-Pp*, but instead of keeping equal proportions of both species by adding fresh bacteria daily to the batch culture, we allowed the bacteria to be carried over during each dilution step. While the bacterial abundances in the media change and reach equilibrium, at the same time and in the same well, competition takes place within the intestine of worms. Migration between these environments takes place in the form of worms feeding from the media and defecating live bacteria into it. We found that despite the strong linkage between environments, *Sm* dominated in the gut yet was present in a minority in the media outside the worm (Fig 3C). Similar differences in competition outcomes between the worm gut and outside media were observed for *Lp-Bs* (Fig S6). These results show that the environmental filtering imposed by the worm intestine can be strong enough to keep an internal bacterial community different from its surrounding environment.

Given the importance of pH to microbial growth and competition (71–75), we tested whether the low pH of the worm gut could cause some of the observed differences between competitive outcomes in the intestine and the media. To this end, we repeated the *Sm-Pp* competition in liquid media at its normal pH 6, and at lower pH 4.5, where the latter approximates the conditions within the nematode intestine (76,77). For this pair of species, lowering the pH of the media was sufficient to alter the pairwise outcome, resulting in a community very similar to that observed in the worm intestine (Fig 3D). Similar results were observed in the pair *Lp-Bs* (Fig S6). We conclude that the acidic pH of the worm gut is an important component of the environmental filtering imposed by this host intestine during community assembly.

### Trio outcomes are predicted by pairwise outcomes

Our results thus far indicate that monoculture colonization and pairwise interactions are both important for the outcomes of pairwise experiments in the *C. elegans* intestine (Fig 3B). However, it remains unclear how these two forces will impact the assembly of more diverse bacterial communities. We therefore constructed three-species communities to extend our analysis. From our eleven bacterial strains, we purposely selected a set of six (*Bc, Lp, Pf, Pv, Ea*, and *Sm*) that covers the different competitive abilities observed. We fed *C. elegans* with the 20 possible three-species combinations of these six species, and measured the bacterial abundances after four days of colonization (Fig S7). To summarize these trio outcomes, we again utilized the mean fractional abundance of each species in all communities where it participates. We found that the population size in monocultures correlates less strongly with this mean fraction in trios (Fig S8, Spearman correlation [r_s_]=.6, *p*=.12) than with the mean fraction in pairwise experiments (Fig 2C). This result indicates that, as community diversity increases, the ability to colonize in monoculture becomes less important.

The mean fractional abundance measured in our experiments agrees with previous reports of the natural *C. elegans* microbiota (35), with our *Enterobacteriaceae, Pseudomonadaceae*, and Firmicute isolates ranking first, second, and third in abundance, respectively. However, more than confirming the potential that each bacterial species has when competing in a natural setting, our bottom-up approach allows us to examine closely many particular microbiomes. For instance, the trio outcome of *Ea-Pf-Sm* is shown in Figure 4A as a stack of bars, and in Figure 4B as a point in a simplex, where the point moves closer to a vertex when that given species increases in abundance. By plotting the measurements of all 20 trios into one simplex (Fig 4C), we observe that most of the cases have one species as highly abundant, yet full exclusion is rare and only accounts for 3 out of the 20 different communities (Fig 4D).

**Figure 4.**
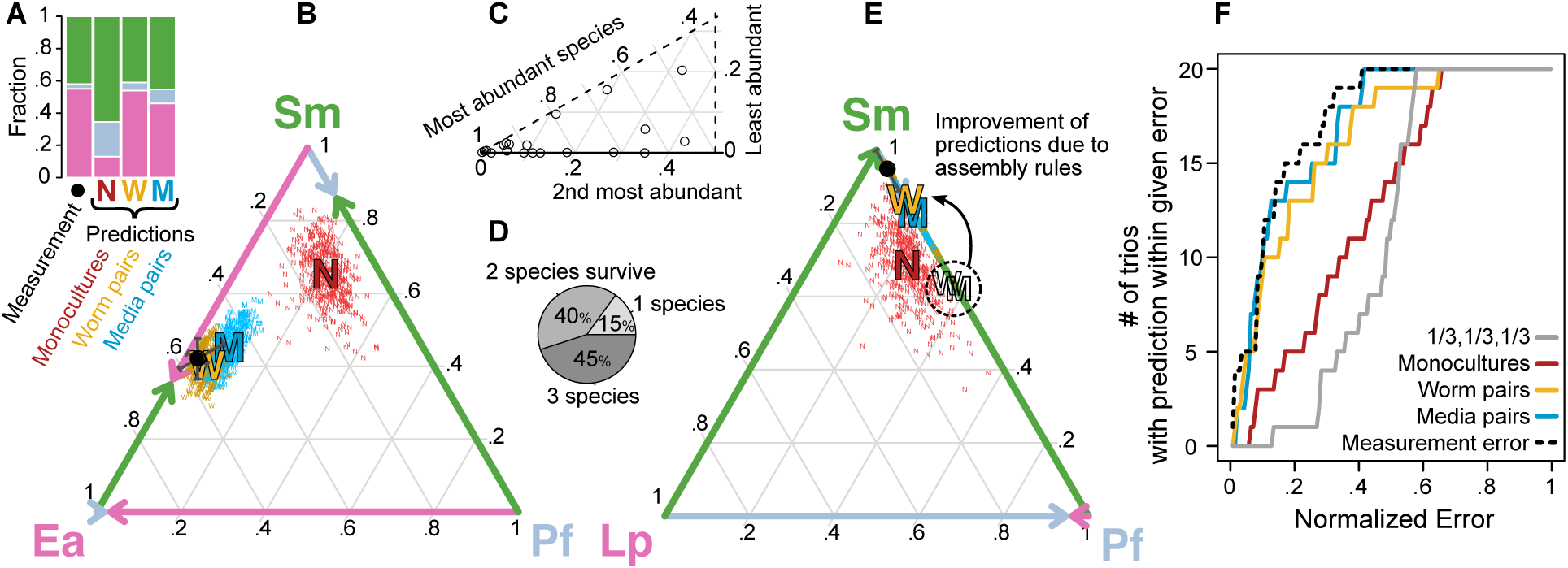
Fractional abundances in three-species communities are well predicted by pairwise outcomes. **(A)** Outcome of trio *Ea-Pf-Sm* in *C. elegans* intestine, together with predictions based on monocultures (each bacterial species reaches its population size in monoculture) or pairwise outcomes (normalized arithmetic mean) in worm or media. **(B)** Simplex representation of trio outcome and predictions in (A), with the edges depicting the pairwise outcomes in the worm intestine. The error bars on measurement are the SEM of 4 biological replicates, and the clouds of small letters around predictions are 400 bootstrap replicates (‘N’s sampling the monoculture data, and ‘W’s and ‘M’s sampling the pairwise data). **(C)** 20 trio outcomes represented in one 6th of a simplex, with the highest, medium, and lowest abundant species taking the left, right and upper vertices, respectively. **(D)** 3, 8, and 9 out of the 20 trios show full competitive exclusion, two- and three-species coexistence, respectively. **(E)** Assembly rules help the quantitative prediction of the trio outcomes when one of the pairwise outcomes is competitive exclusion. **(F)** Cumulative distribution of error of predictions. Error calculated as the linear distance between prediction and measurement in the simplex. The distances are normalized by the maximal distance, 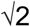. The dashed line represents the mean distance between the measured mean and the biological replicates of each trio.

As with the pairwise outcomes, the mean abundance of *Ea* is the least predictable based on its monoculture colonization (Fig S8), which suggests that the pairwise interactions favoring the abundance of *Ea* in co-cultures might favor *Ea* further in larger communities. To determine if pairwise interactions are the main drivers of trio outcomes, we made predictions of all trios based on either monocultures, or pairwise outcomes. To make quantitative predictions of trios based on monocultures, we extended the null expectation described earlier by assuming that all species will reach their monoculture population sizes (‘N’s in Fig 4 and S7). This null expectation based on monoculture colonization achieves poor results at predicting trio communities, for its mean error of 35.7% is just slightly better than the 43.8% mean error of an uninformed ‘⅓, ⅓, ⅓’ prediction (Fig 4F). Hence, although monoculture colonization predicts pairwise outcomes with a lower mean error of 20% (Fig S3), monocultures become inadequate at predicting three-species microbiomes, highlighting again the importance of interspecies interactions.

Alternatively, we can predict trios based on pairwise outcomes by taking the arithmetic mean of each species’ abundance in the co-culture experiments against the other two species (here, a normalization factor of 2/3 is needed for the fractions of the three species to add to one). This *normalized arithmetic mean* prediction, applied to the pairwise outcomes obtained in the worm intestine or the *in vitro* liquid media (‘W’s and ‘M’s in Fig 4A and B, respectively), quantitatively predicts some trios with high accuracy, and reaches a lower mean error of 26% (Fig S9). However, this simple prediction is prone to error when one of the pairwise outcomes is competitive exclusion, such as *Sm-Lp* in the *Sm-Lp-Pf* trio (Fig 4E, empty letters). Fortunately, a recently proposed ‘assembly rule’ (69) is capable of adjusting these cases by simply removing a bacterial species from the trio prediction when it cannot survive all pairwise competitions (Fig 4E and S7). After the application of these assembly rules, the mean error of the predictions based on worm or media pairwise outcomes, 18.7 and 15.7%, comes close to the expected biological noise in trio outcomes, 13.3% (Fig 4F). The fact that pairwise outcomes can properly predict trio outcomes indicates that interactions between pairs of bacterial species are an important force in determining the outcome of multispecies communities and suggests that indirect interactions are uncommon or weak.

## Discussion

Here we characterized the outcomes of all co-culture experiments among 11 bacterial species within the gut of the worm *C. elegans*. We find that monoculture population size in the worm intestine correlates well with competitive performance in this environment, suggesting that the ability to colonize and proliferate in the host is an important determinant of success. However, we find that the outcomes of pairwise competition, but not single-species colonization ability alone, predict the outcomes of three-way competition, indicating that pairwise competitive interactions between microbes are an important driver of the assembly of more complex multispecies communities.

We found that most interactions among our set of species are competitive, and none of the 55 pairs that we measured displayed a strong mutualism in which both species benefited. Despite this prevalence of competitive interactions, we also found that coexistence between species was very common, indicating that the competition is relatively weak and only rarely leads to competitive exclusion. Interestingly, we do not observe any cases of strictly non-transitive interactions (rock-paper-scissors) (66) among any of the 165 possible trios, questioning once more the practical significance of this mechanism at stimulating coexistence and diversity in multispecies communities (63). However, the 11 species that we studied here are laboratory isolates rather than a naturally occurring community, which may have evolved to co-occupy their particular environment (78). Such communities could be enriched for mutualistic or rock-paper-scissors interactions; future work will be necessary to determine whether naturally occurring communities have significantly different network properties, although recent work suggests that non-transitive interactions are not common in natural microbial communities (79).

Although we observed a range of different outcomes, from coexistence to competitive exclusion, we have not extensively explored the mechanistic basis behind them. These outcomes could arise from simple resource competition, including competition for the limited space available within the *C. elegans* gut, or there could be more explicit forms of antagonism such as toxin production. In cases of coexistence, spatial partitioning within the host could play an important role; the role of spatial structure was recently found to be important in determining the competitive outcome of interspecies competition within the gut of the zebrafish (17). Further work will be required to elucidate these spatial dynamics within the worm.

Our results provide additional support for the importance of host-determined environmental factors, specifically pH, in shaping the microbiome. The low pH of the gut environment is thought to be a critical factor in host-microbe interactions, and recent work has explicitly demonstrated the importance of pH in modulating the interactions between microbes and determining the structure of synthetic and natural communities (71–73). Consistent with these results, we observed that reducing the pH of a liquid medium to simulate the host intestine could alter the outcomes of competition between species and substantially reduce the difference between *in vitro* media and *in vivo* gut.

The conserved competitive hierarchy among our set of microbial species inside and outside the worm suggest that similar traits are required for success in both environments. Notably, these experiments were carried out in an immune-deficient host strain (AU37) which is defective in recognition of and response to microbial colonization (80). It remains to be seen whether the more stringent environment of an immune-capable host would produce larger differences between *in vitro* and *in vivo* experiments. In future work, it will therefore be interesting to determine the extent to which modifying the properties of the host environment alters the filtration imposed by that environment and the interactions between colonizing microbial strains.

The work presented here focuses on population averages rather than the composition of bacterial communities within individual worms. Recent results from our group have demonstrated that variation between individuals can be informative - when colonization is very slow, there can be an extreme bottleneck during colonization of the worm gut that leads to marked heterogeneity between the gut communities in different worms (81). Similar results have recently been found in *Drosophila* (82), indicating that stochastic effects during colonization may be important in a wide range of host species. Under the feeding densities used in these experiments we expect more uniform colonization, but it will be important for future studies to determine the role of stochasticity and priority effects during assembly of multispecies communities within the host.

Our results are based on a single time point after multiple days of colonization, at which point we expect any initial transients in population dynamics to have died out. This experimental choice allows us to focus on community composition after species have had time to interact, but also precludes the study of the early colonization dynamics. *C. elegans* in the wild is born sterile and is then colonized by the complex microbial communities present within the soil and rotting organic matter (83). This is similar to the microbial successions that have been observed in other contexts, such as in the digestive tract of human infants, *in situ* and *in vitro* cheese rind communities, enrichments of seawater in chitin particles, and *in vitro* enrichments of leaf communities (84–87). It would be fascinating to determine whether similar succession dynamics take place during early colonization of the worm intestine.

In this study, we have fed worms with monocultures, pairs, and trios of bacteria from a set of naive soil isolates to determine the role of interspecies interactions in the assembly of host-associated microbial communities. These results add to our understanding of how interactions between pairs of bacterial species can inform our understanding of more complex bacterial communities. Our results show the promise of experimental bottom-up microbial ecology as a tool for understanding the dynamics of bacterial gut communities in a simple model organism, providing insight into the forces that shape and control the structure of microbiomes.

## Methods

### Nematode culture

Nematodes were grown, maintained, and manipulated with standard techniques (88). The *C. elegans* strain AU37 (*glp-4(bn2)* I; *sek-1(km4)* X) was used for all experiments. Due to the *glp-4* mutation, AU37 does not develop gonads and is therefore sterile when raised at room temperature (23-25°C). Sterility ensured that all worms in a given experiment were of the same age and had the same life history. The mutation on the *sek-1* gene, part of the p38 MAPK signalling pathway (80), decreases immune function and makes AU37 more susceptible to bacterial colonization (41). The ensuing larger intestinal bacterial communities allowed better quantification of fractional abundances. Worm strains were provided by the Caenorhabditis Genetic Center, which is funded by NIH Office of Research Infrastructure Programs (P40 OD010440).

L1 animals of the same age were obtained using the standard egg-extraction protocol (88). Starved larvae were then transferred to NGM plates with lawns of *E. coli* OP50, and after 2 days of feeding at room temperature, a synchronous adult population of *C. elegans* was obtained. Worms were then transferred to S-medium (88) + 100 µg/mL gentamicin + 5X heat-killed OP50 for 24 hours to kill any bacteria inhabiting the intestine, resulting in germ-free worms. Heat-killed OP50 was used to trigger feeding behaviour in the worms. The adult worms were washed via sucrose flotation to remove debris before bacterial colonization.

### Bacteria

*Bacillus subtilis* (ATCC 23857), *Enterobacter aerogenes* (ATCC 13048), *Lactobacillus plantarum* (ATCC 8014), *Pseudomonas aurantiaca* (*Pseudomonas chlororaphis* subsp. *aurantiaca*) (ATCC 33663), *Pseudomonas citronellolis* (ATCC 13674), *Pseudomonas fluorescens* (ATCC 13525), *Pseudomonas putida* (ATCC 12633), *Pseudomonas veronii* (ATCC 700474), *Serratia marcescens* (ATCC 13880) were obtained from ATCC. *Bacillus cereus* was obtained from Ward’s Scientific Catalog. *Escherichia coli* MC4100 (CGSC #6152) was obtained from the *E. coli* Genetic Stock Center.

Bacterial strains were grown for 24hrs at 30°C in individual culture tubes with 2 mL of LB (Difco). To construct cultures to feed *C. elegans*, strains were centrifuged 1 minute at 7K RCF to pellet, washed once in S-medium, and then resuspended in 1% v/v Axenic Medium diluted in S-medium (1%AXN). 100 ml of 100% AXN were prepared by autoclaving 3g yeast extract and 3g soy peptone (Bacto) in 90 ml water, and subsequently adding 1g dextrose, 200µl of 5 mg/ml cholesterol in ethanol, and 10 ml of .5% w/v hemoglobin in 1 mM NaOH. To standardize the bacterial densities used to feed *C. elegans*, the bacterial cultures were diluted to ∼10^8^ CFU/ml based on previously measured CFU counts.

### Bacterial colonization of C. elegans

Germ-free adult worms were resuspended in 1%AXN to a concentration of ∼1000 worms/mL. Aliquots of 120µL (∼100 worms) were laid into 96-deep-well culture plates (1 mL well volume, Eppendorf). 15µL of each bacterial suspension were added to the corresponding wells in a matrix-like format (11 species x 11 species), leaving a final bacterial concentration of ∼2×10^7^ CFU/ml, and a 150µl volume for monoculture and co-culture experiments. Extra replicates of monoculture colonization were done to complement the diagonals of the matrices containing the co-culture experiments. For trio competitions, 3 bacterial species were mixed evenly to reach a concentration of ∼3×10^7^ CFU/ml. Plates were covered with a Breathe-Easy sealing membrane and incubated with shaking at 400 RPM at 25°C. To maintain the bacterial concentrations constant throughout the 4-day feeding period, every day the worm samples were washed and the bacteria was replenished. Samples were washed with a VIAFLO 96 by adding 500µl of M9 Worm Buffer (WB) + 0.1% v/v Triton X-100 (Tx), pipetting 10 times, and removing the supernatant after worms precipitated. The samples were then transferred to new 96-deep-well plates to leave behind possible biofilms, and then washed in the same way two more times with 1%AXN. Bacterial suspensions from new culture tubes were added as previously described.

### Mechanical disruption of worms

The worm samples were washed to remove most external bacteria with the previously described protocol. Samples were then incubated in 100µL S-medium + 2X heat-killed OP50 at 25°C for one hour to allow the worms to evacuate any non-adhered bacterial cells from the intestine. Worms were then rinsed twice with WB + 0.1%Tx, cooled down 15 min at 4°C to stop peristalsis, and bleached for 6 minutes at 4°C with 100µL WB + .2% v/v commercial bleach. Worms were then rinsed three times with cold WB + 0.1%Tx to remove bleach.

For manual disruption, each worm sample was transferred to a small petri dish (6 cm) with 3mL of WB + 1%Tx (the high concentration of surfactant facilitates the upcoming grinding). To guarantee the background media was fully clean, 20µl of the supernatant in each petri dish were collected, serially diluted, and laid in agar plates (see below). Then, a batch of ∼20 worms was collected by pipetting 200µl from each petri dish into a 0.5 mL microcentrifuge tube (Kimble Kontes), and the exact number of collected worms was recorded. The volume of each microtube was then reduced to 20µl and the worms were grinded for one minute with a motorized pestle (Kimble Kontes Pellet Pestle, Fisher Scientific). After disruption, tubes were centrifuged two minutes at 7K RCF to collect all material, and the resulting pellet was resuspended in 180µL WB (final volume 200µL) before transfer to 96-well plates for serial dilution in 1X phosphate-buffered saline (PBS).

For 96-deep-well plate disruption, rinsed worms were treated for 20 minutes with 100µL of a solution of SDS + DTT (WB + 0.25% v/v sodium dodecyl sulfate + 3% v/v dithiothreitol [1M, freshly mixed in water], chemicals from Sigma Aldrich) to partially disrupt the cuticle, and then washed twice in WB + 0.1%Tx. A deep-well plate (2 mL square well plate, Axygen) was prepared by adding ∼0.2g of a sterile grit (36-mesh silicon carbide, Kramer Industries, autoclaved) to each well. Same as in the manual disruption, the worm samples were transferred to a small petri dish and supernatant samples were taken. Then, batches of ∼20 worms were pipetted into individual wells of the Axygen plate. The plate was covered with parafilm and kept at 4°C for one hour prior to disruption to reduce heat damage to bacteria. Parafilmed plates were capped with square silicon sealing mats (AxyMat) and disrupted by shaking at high-speed (Retsch MM400, 30 hz for 3 minutes). Plates were then centrifuged at 3K RCF for 2 minutes to collect all material, resuspended by pipetting, and transferred to 96-well plates for serial dilution in PBS.

### Measurement of bacterial fractional abundances

The undiluted worm digests and supernatants, together with the serial dilutions 10^-1^, 10^-2^, and 10^-3^ of each sample, were plated onto Nutrient Agar (3g yeast extract, 5g peptone, and 15g of agar [Bacto] in one liter of water). The samples were incubated at room temperature for two days to allow distinct colony morphologies to develop. Colonies in all distinguishable dilutions were counted afterwards.

Pairwise and trio outcomes were categorized as coexistence if the rare species was present at an average abundance of more than 2%. This threshold is just above our usual limit of detection of ∼1%, which is inversely proportional to the number of colonies counted (∼100). The pair *Pf-Ea* (1.7%-98.3%) was defined as coexisting since we could reassure the presence of *Pf* with more than one biological replicate.

### Hierarchy scores and generation of random matrices

Utilizing the 11×11 matrix with the fractional abundances in co-culture experiments, the hierarchy score is calculated by: 1) Ordering its rows and columns ascendingly based on the mean fractional abundance; and 2) Taking the mean value of the half-matrix under the diagonal. In a perfectly hierarchical matrix, each competitor will drive to extinction every other species with a lower rank, reaching a hierarchy score of 1. Random matrices were generated to calculate the significance of the observed high hierarchy score. We conserved the distribution of fractional abundances by: 1) Sampling with replacement 55 values of the original matrix; 2) Assigning these random fractional abundances to the lower triangle of a new matrix; and 3) Assigning to the upper triangle of the matrix the values of 1-transpose. For each random matrix generated, a new hierarchy score is calculated as previously described.

### Null expectation of fractional abundances

The null expectation based on monocultures is obtained by averaging the fractional abundances in all possible combinations of monoculture information. Since we have ∼10 replicates for each species colonizing *C. elegans* in monoculture, the null expectation for the pairwise and trio outcomes is the mean of ∼100 and ∼1000 different combinations of monoculture data, respectively. The SEM is calculated by dividing the standard deviation of all combinations by sqrt(least number of monoculture replicates). These mean and SEM can also be obtained by bootstrapping (sampling with replacement) the monoculture data.

### Co-culture experiments in vitro

Utilizing the same S-medium + 1%AXN, 96-deep-well culture plates, 150µl volume per sample, and the same matrix format used to colonize *C. elegans*, pairs of bacterial species were mixed at a concentration of 10^5^ CFU/ml each. We allowed the bacterial relative abundances’ to equilibrate with seven growth-dilution cycles, where the bacteria are diluted 100-fold into fresh media each day. As previously described, we quantified the bacterial abundances by plating into agar.

### Phylogeny reconstruction

Sequences of the full 16S rRNA gene were obtained from NCBI. *Sulfolobus solfataricus*, a thermophilic archaea, was used as an outgroup species to root the tree. Clustal X with default parameters was used to align the sequences (89). PhyML-SMS with default parameters was used to select GTR+G+I as the best model and to infer the tree (90).

