## Supplemental Figures for "Interspecies bacterial competition determines community assembly in the *C. elegans* intestine"

### Supplementary Figures

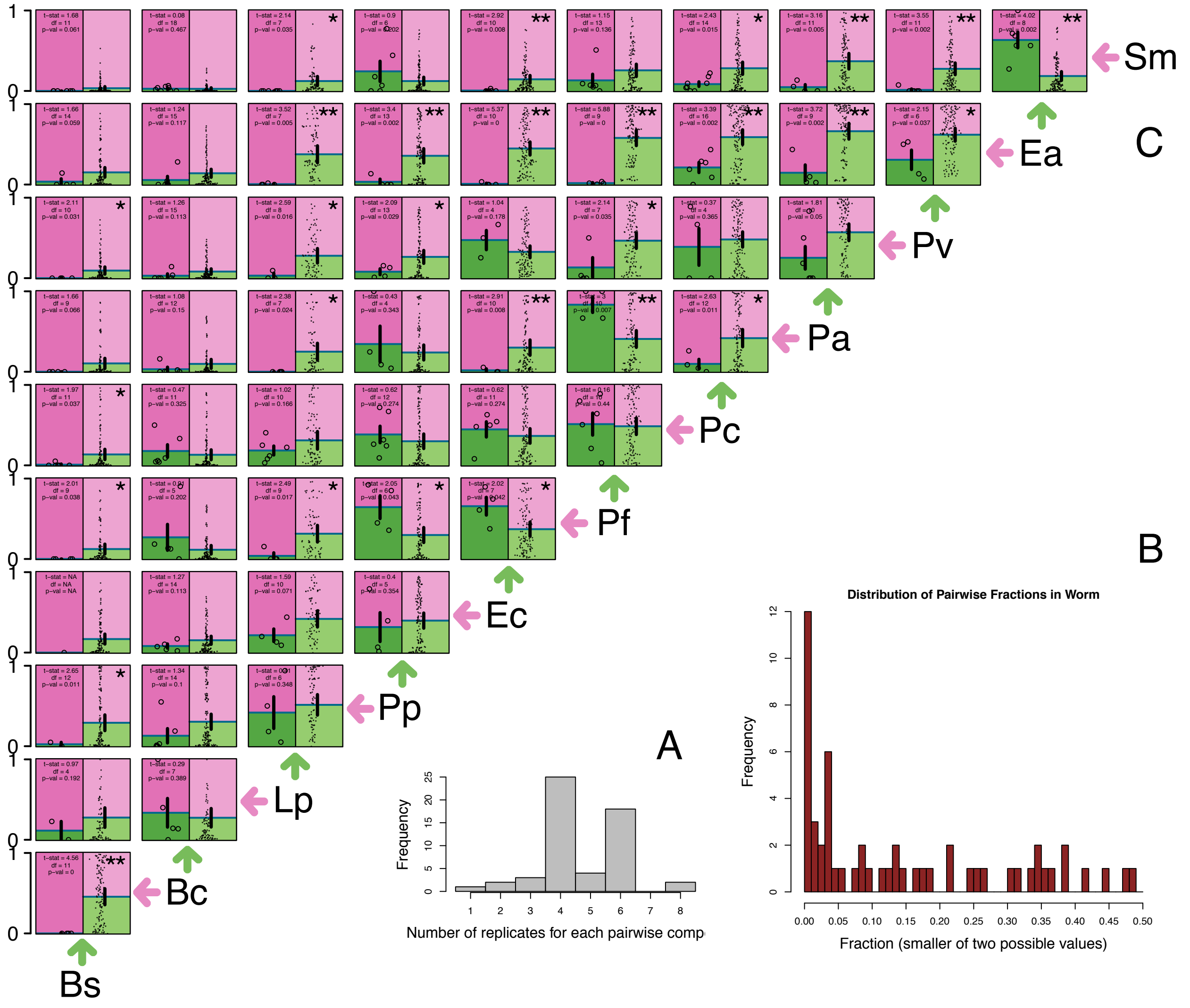


**Figure S1. Replicates of pairwise experiments in *C. elegans* intestine. (A)** From the (11 choose 2)=55 different pairs of bacteria colonizing the *C. elegans gut*, most conditions had four or more biological replicates. Each replicate involves the colonization and disruption of an independent set of worms. No more than two replicates were done in the same date. *Bs-Lp* and *Bs-Pp* had 2 replicates, and *Bs-Ec* had one replicate; several attempts to quantify these and other pairs failed due to low amounts of worms at the end of the experiment and/or zero CFU counts. **(B)** Histogram of the fractional abundances in co-culture experiments. From a pair of bacteria reaching fractions 57%-43%, only the lower quantity, 43%, was plotted. **(C)** Same as Fig 2A (Fractional abundances in the left, and null expectation based on monoculture colonization on the right. Error bars as standard error of the mean), but including the replicates of each pairwise experiment as points in left panels, and the fractions obtained from all combinations of monoculture population sizes as points in right panels.

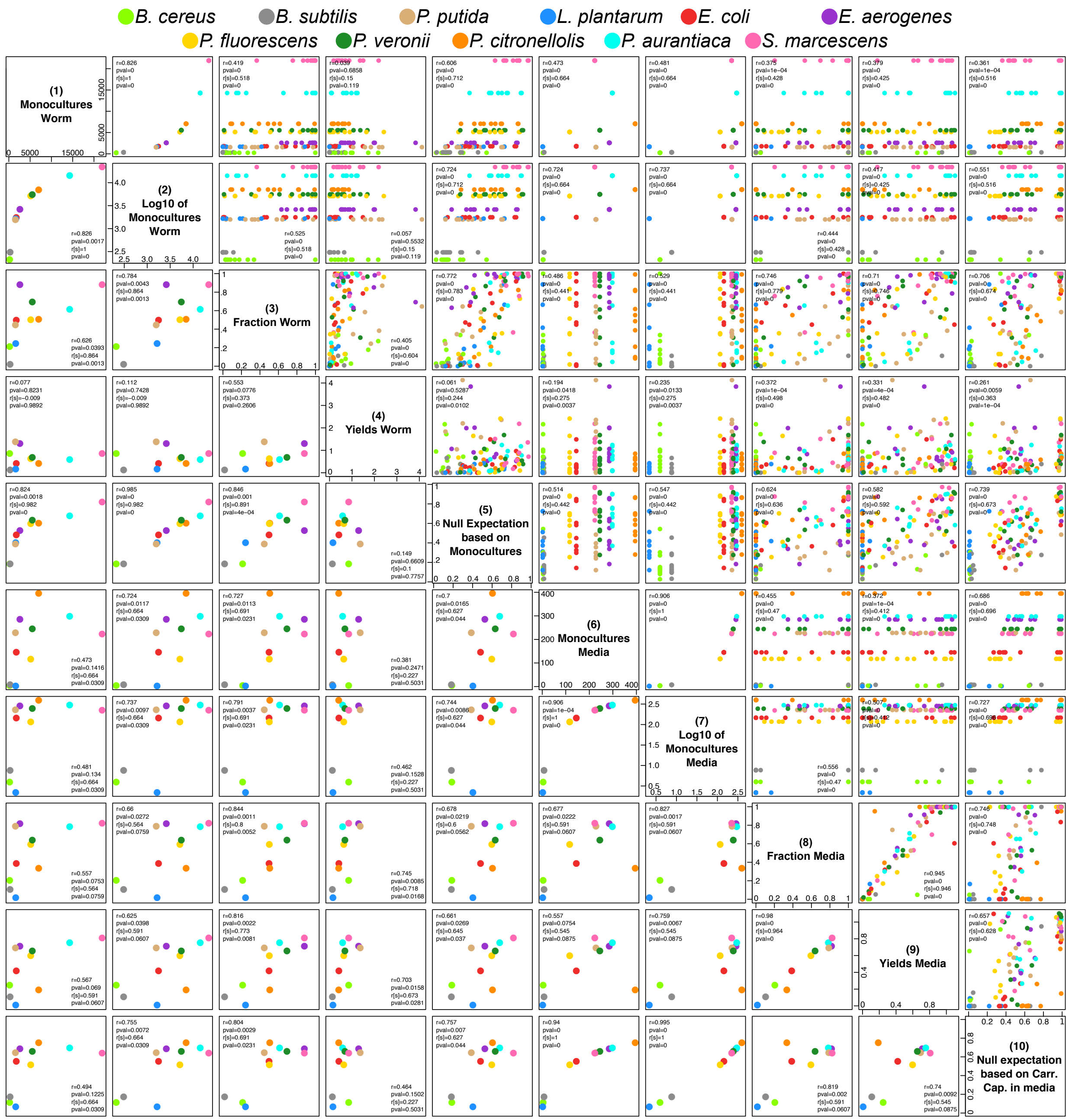


**Figure S2. Pearson and Spearman correlations between different ecological metrics.** 11 Bacterial species (names at the top) were grown in monoculture and co-culture in the *C. elegans* intestine and *in vitro* liquid media. From these experiments, we calculated: **(1)** the colony forming units (CFU) per worm in monoculture colonization of the gut, and **(2)** their log10 transformation; **(3)** the bacterial fractional abundances in co-culture experiments in the worm gut; **(4)** the *relative yields* in the worm as the population size of a species in co-culture experiments divided by its population size in monoculture; **(5)** a null expectation for the fractional abundances in the worm gut assuming that each species is able to reach the carrying capacity that was measured in monoculture colonization; and **(6 to 10)** these same metrics but *in vitro* liquid media (the monoculture carrying capacity in media is the number of CFU in 10µl of 1%AXN). Each subpanel contains the comparison between 2 different metrics. The subpanels above the diagonal display all the pairwise combinations, while the subpanels below the diagonal display the average metric across all competitors (i.e. a point in [row 3, column 8] is the fraction of the bacteria denoted by the color in one single co-culture experiment, while a point in [8,3] is the average fraction of that bacteria in ten co-culture experiments). The *p*-values for the Pearson correlations, ‘r’, and Spearman correlations, ‘r[s]’, were calculated with *cor.test* in *R* using a two sided alternative hypothesis.

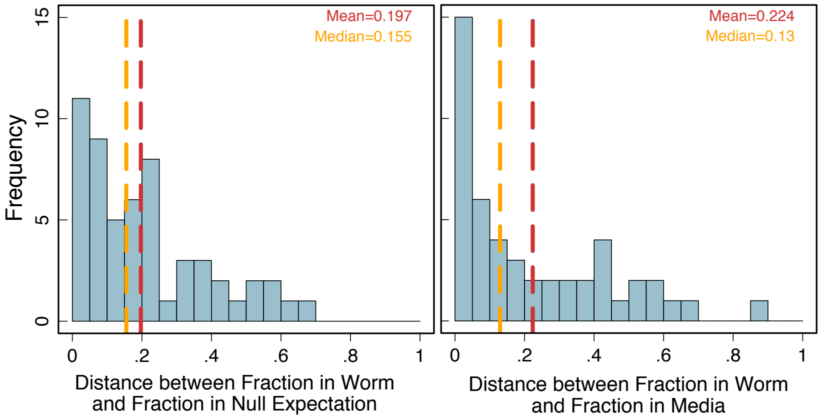


**Figure S3. Differences between pairwise outcomes in worm intestine and null expectation and *in vitro* liquid media. (Left)** Distribution of differences between the co-culture experiments in the worm intestine and a null expectation based on the monoculture colonization of the worm intestine. The null expectation assumes a non-interactive scenario where each bacterial species is capable of reaching its own measured monoculture population size. (This is the histogram of the distances between the left and right panels of Figure 2A.) **(Right)** Distribution of differences between co-culture experiments in the worm intestine and co-culture experiments *in vitro* liquid media. (This is the histogram of the distances between the blue circles and the identity line in Figure 3B.) The similitude in mean and median between these two distributions shows how monoculture colonization and pairwise outcomes measured in a different environment can predict with similar accuracy the pairwise outcomes in the worm intestine.

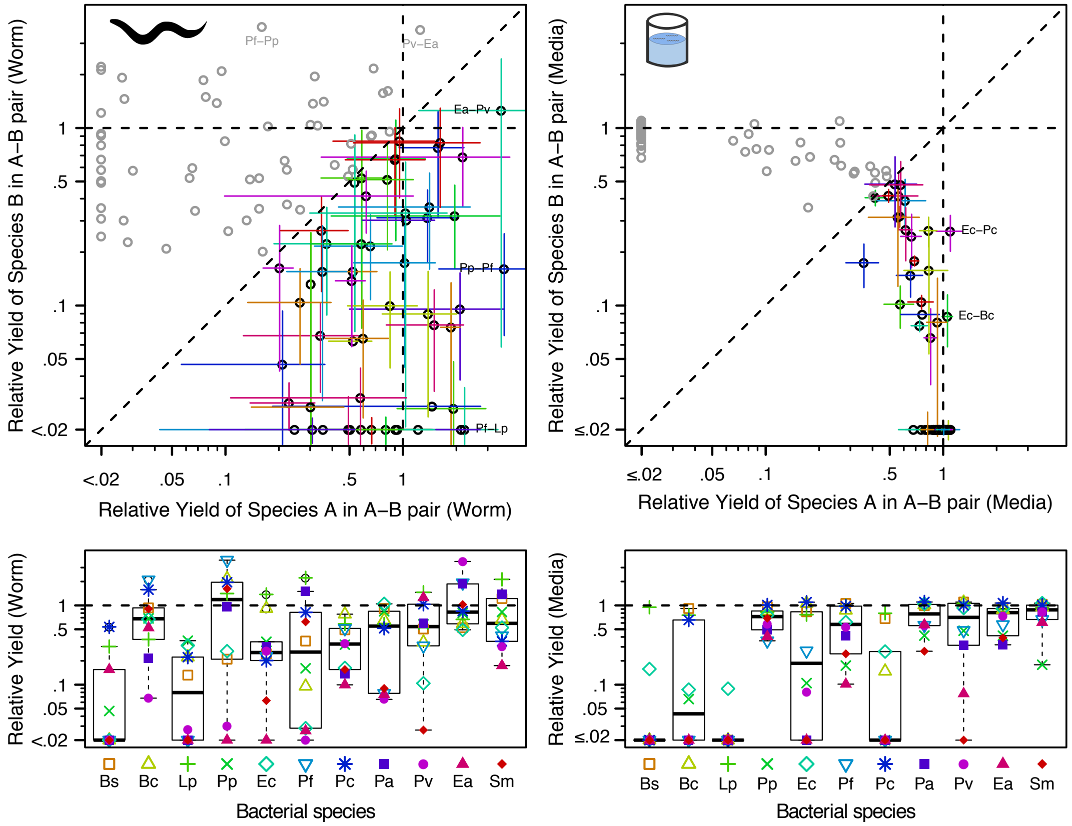


**Figure S4. Relative yields in co-culture experiments in the worm intestine and *in vitro* liquid media reveal competitive interactions.** The relative yield of a bacterial strain is calculated by dividing its population size in a pairwise experiment (average CFUs per worm across 2 to 8 replicates [Fig S1], or average CFUs per µl of media across 6 replicates [Fig S5]) by its monoculture population size. In all four subpanels, each point represents one pairwise combination. **Upper panels:** Each point displays the relative yield of both co-cultured species. Error bars as the SEM. Some points were labeled, the error bars were colored, and dotted lines were added solely to improve clarity. The points above the diagonal are equivalent to the points below the diagonal, and their error bars would be equivalent as well. **Lower panels:** Box plots, where each point displays the relative yield of the species denoted at the bottom when co-cultured with the species denoted by the colored symbol. 19.1% (21/110) and 13.6% (15/110) of the relative yields measured in the worm intestine and liquid media were above 1, respectively, denoting an enrichment for competitive interactions. The points above one are heavily influenced by unique replicates of co-culture experiments with an abnormally large community size.

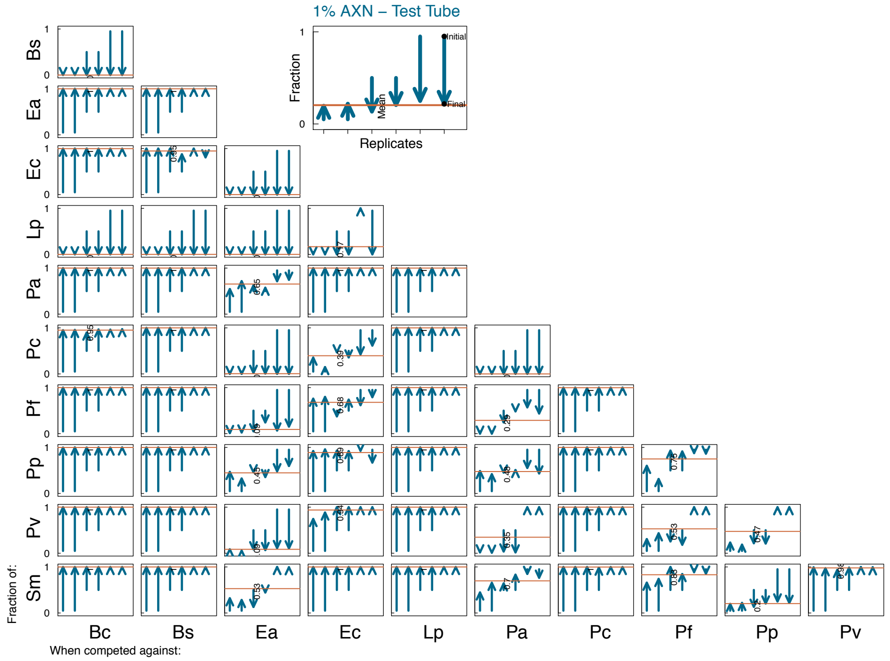


**Figure S5. Fractional abundances in co-culture experiments *in vitro* liquid media.** Pairwise experiments in the background liquid medium (1%AXN), with each subpanel representing one pairwise combination, and each arrow representing a replicate. The starting fraction of each replicate is denoted by the beginning of the arrow, and the measured fraction after 7 cycles of daily dilution is displayed by the end of the arrow. The orange lines are the means of the replicates.

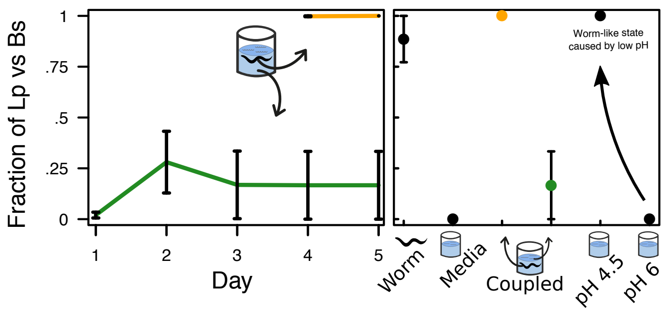


**Figure S6. Low pH in worm intestine sustains a different microbial composition than the liquid media. (Left)** Fraction of *Lp* in worm and media when environments are coupled by migration. Both environments reach their different equilibrium points in the same test tube. **(Right)** An acidic version of the media resembling the average pH of the worm (4.5) shifts back the pairwise outcome of *Sm-Pp* to a worm-like state.

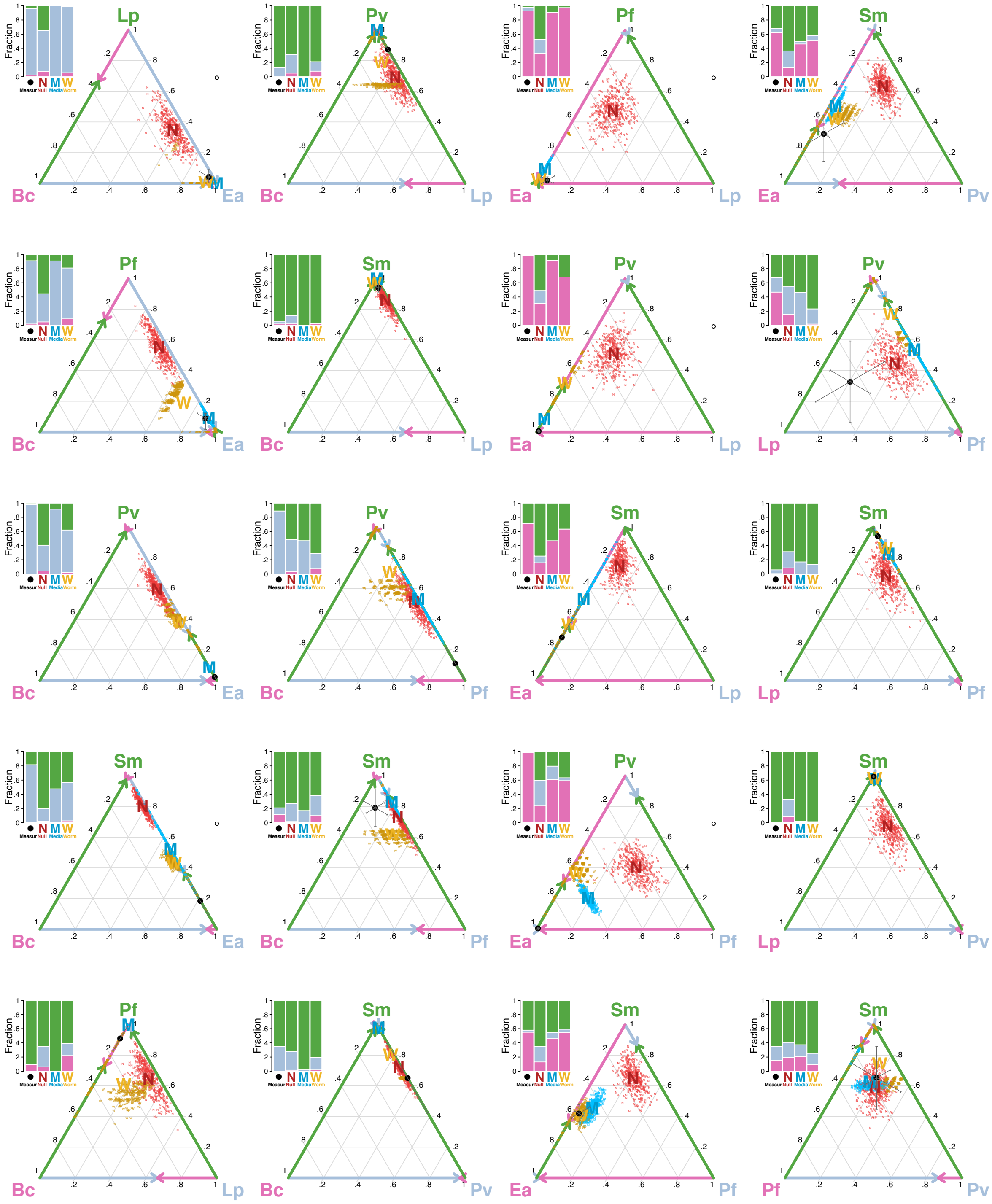


**Figure S7. Measurements of 20 three-species microbiomes in the *C. elegans* intestine, and predictions of them based on one- and two-species microbiomes.** Each set of bar graphs and the triangle (simplex representation) to the right displays the measured fractional abundances and predictions of one three-species community. The two-letter codes on the vertices of each simplex denote the species in that trio. The arrows in the edges of the simplexes depict the pairwise outcomes in the worm intestine.

**Red ‘N’s**: predictions based on monocultures, where each bacterial species reaches its population size in monoculture. **Blue ‘M’s**: predictions based on pairwise outcomes *in vitro* liquid media (normalized arithmetic mean after applying assembly rules). **Golden ‘W’s**: predictions based on pairwise outcomes in worm intestine. The error bars on measurement are the SEM of 4 biological replicates, and the clouds of small letters around predictions are 400 bootstrap replicates (‘N’s sampling the monoculture data, and ‘W’s and ‘M’s sampling the pairwise data).

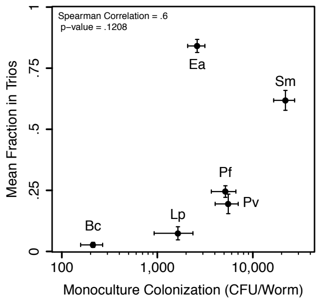


**Figure S8. The mean fractional abundance in three-species microbiomes correlates mildly with monoculture population size.** X-axis as the average population size of 8 or more biological replicates of monoculture colonization, and its error bars as the SEM. Y-axis as the mean fractional abundance across all possible three-species combinations of these six bacterial species, and its error bars as the propagated error from the SEM of the four replicates of each trio.

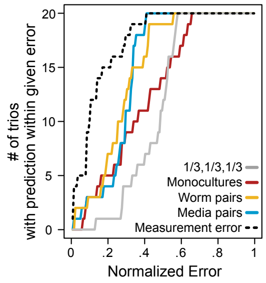


**Figure S9. Accuracy of predictions of three-species communities without *assembly rules*.** Cumulative distribution of the error of three different predictions. The prediction of three species communities based on monocultures assumes that all species will reach their monoculture population sizes, and the predictions based on pairwise outcomes are the arithmetic mean of each species’ abundance in the co-culture experiments against the other two species. The error of the predictions based on pairs can be reduced by removing a bacterial species from the trio prediction when it cannot survive all pairwise competitions—assembly rules (Fig 5F). Errors calculated as the linear distance between prediction and measurement in a simplex (normalized by the maximal distance, sqrt[2]). The dashed line represents the mean distance between the measured mean and the biological replicates of each trio, and it is an upper bound for the error. The gray line is the error of an uninformed prediction of ‘1/3, 1/3, 1/3’, and it is a lower bound for the error.
